# Sustained activity in a descending neuron is associated with flight saccades in *Drosophila*

**DOI:** 10.1101/2024.08.22.608914

**Authors:** Elhanan Buchsbaum, Bettina Schnell

## Abstract

Flies perform rapid turns termed saccades to change direction during flight. Evasive turns can be elicited by looming stimuli mimicking an approaching object such as a potential predator. Whereas projection neurons of the optic lobes responsive to looming stimuli have been well described, how this information is transmitted to the motor system to elicit a saccade is not well understood. Here we describe activity of the descending neuron DNp03 in *Drosophila*, which receives direct input from looming-sensitive visual interneurons and projects to wing motor areas within the ventral nerve cord. Whole-cell patch-clamp recordings from this neuron during head-fixed flight confirm that DNp03 is responsive to looming stimuli on the side ipsilateral to its dendrites. In addition, activity of this neuron is state-dependent as looming stimuli only elicit spikes during flight and not rest. The behavior in response to the looming stimulus is variable, which allowed us to study how activity of DNp03 relates to the execution of a saccade. Our analysis revealed that sustained activity in DNp03, persisting even after the visual stimulus ended, was the strongest predictor of saccade execution, therefore reflecting the behavioral decision of the fly to respond to the stimulus.

## Introduction

Evading from an approaching predator is essential for the survival of small animals. Approaching objects are detected as visual looming meaning a rapid expansion of a stimulus as perceived by the eye. Visual looming triggers defensive responses in a wide variety of organisms ranging from mice to fish to insects ^1–4^ and can elicit different behaviors such as freezing, flight, turning or take-off. Fast take-off responses in fruit flies have e.g. been shown to increase their survival when faced with potential predators and their neuronal underpinnings are well described ^5,6^. The fly’s visual system is therefore well suited to detect stimuli that increase rapidly in size on their retina as evidenced by the many visual projection neurons within the lobula complex of *Drosophila*, a part of the optic lobes, that encode looming stimuli ^7–10^. These neurons have different receptive fields and project to the optic glomeruli in the central brain, where they make direct synapses with descending neurons such as the giant fiber, which is known to mediate escape jumps in response to looming stimuli in sitting flies ^6,11^. In contrast, a looming stimulus during flight typically elicits a rapid turning response called a saccade ^12^. During a saccade a fly can change its flight direction within about 50 ms reaching angular velocities of over 1000°/s ^13^. However, it is still unclear, how these maneuvers are controlled by the nervous system of the fly. Saccades can not only be initiated in response to an imminent collision or attack, but also occur spontaneously ^14–17^. In flies, saccades can be studied using a head-fixed preparation, where they can be measured as rapid changes in the difference between left and right wing stroke amplitude (L-R WSA), even when flies cannot actually turn ^18,19^. Previous studies using this preparation have shown that two descending neurons (DNs), DNaX and DNb01, exhibit activity correlated with both spontaneous and looming-elicited turns ^20,21^. Moreover, artificial activation of these neurons is sufficient to trigger fictive saccades. However, these two types of neurons do not receive direct input from visual interneurons ^21–24^. In contrast, other descending neurons have arborizations within optic glomeruli and were implicated in the execution of behavioral responses elicited by visual looming, such as escape jumps and landing ^25–27^. One of them, DNp06 was shown to contribute to the execution of saccades ^28^, but it responds to looming stimuli coming from both sides ^29^ and genetic manipulation experiments lead to only subtle changes in behavior, suggesting that other descending neurons play a major role in controlling saccades.

DNp03 constitutes a pair of descending neurons with cell bodies on the posterior side of the brain. It receives a significant fraction of its input from looming-sensitive LPLC1 and LC4 neurons on its dendritic tree spanning multiple optical glomeruli ^26,27^. It then crosses the midline and makes synapses with other descending neurons similar to DNaX and DNb01 ^22^. Within the VNC, it makes direct synapse with wing motor neurons involved in steering during flight ^30^. Its connectivity therefore makes it a good candidate for controlling flight saccades. However, its response properties have not yet been described. We therefore performed whole cell patch-clamp recordings from DNp03 in a head-fixed preparation ^19^, which allowed us to study its activity during flight. We show that DNp03 is responsive to visual looming on its ipsilateral side in a behavioral-state dependent fashion. The variability in saccade execution in our preparation allows us to examine the correlation between DNp03 activity and turning behavior, providing insights into the neural processes underlying differing behavioral responses to identical visual stimuli. Our analysis revealed that sustained activity in DNp03, persisting even after the visual stimulus ended, was the strongest predictor of saccade execution. However, DNp03 activity alone could not fully account for the observed variability in behavioral responses. Together with results from optogenetic activation experiments during free flight, this result suggest an important, but not exclusive role for DNp03 in controlling saccades.

## Results

### Responses of DNp03 to visual stimuli

Based on its connectivity with looming-sensitive visual interneurons, the descending neuron DNp03 seemed a good candidate for controlling saccades during flight. Using a sparse split-Gal4 driver line ^27^, we expressed GFP in DNp03 (Fig. 1A) to target it for whole-cell patch clamp recordings using a preparation in which the wings remain free to move so that we can measure turning as changes in L-R WSA (Fig. 1B) ^19,31^. We only recorded from the cell body of the neuron on the right side of the brain. Stimuli presented in the right visual field are therefore ipsilateral with respect to cell body position, and stimuli in the left visual field contralateral. The average membrane potential of all recorded cells included in the analysis after correcting for the liquid junction potential was -58 mV. An example of the expression pattern within the brain as well as the recorded cell filled with Biocytin is shown in Fig. 1A.

**Figure 1.**
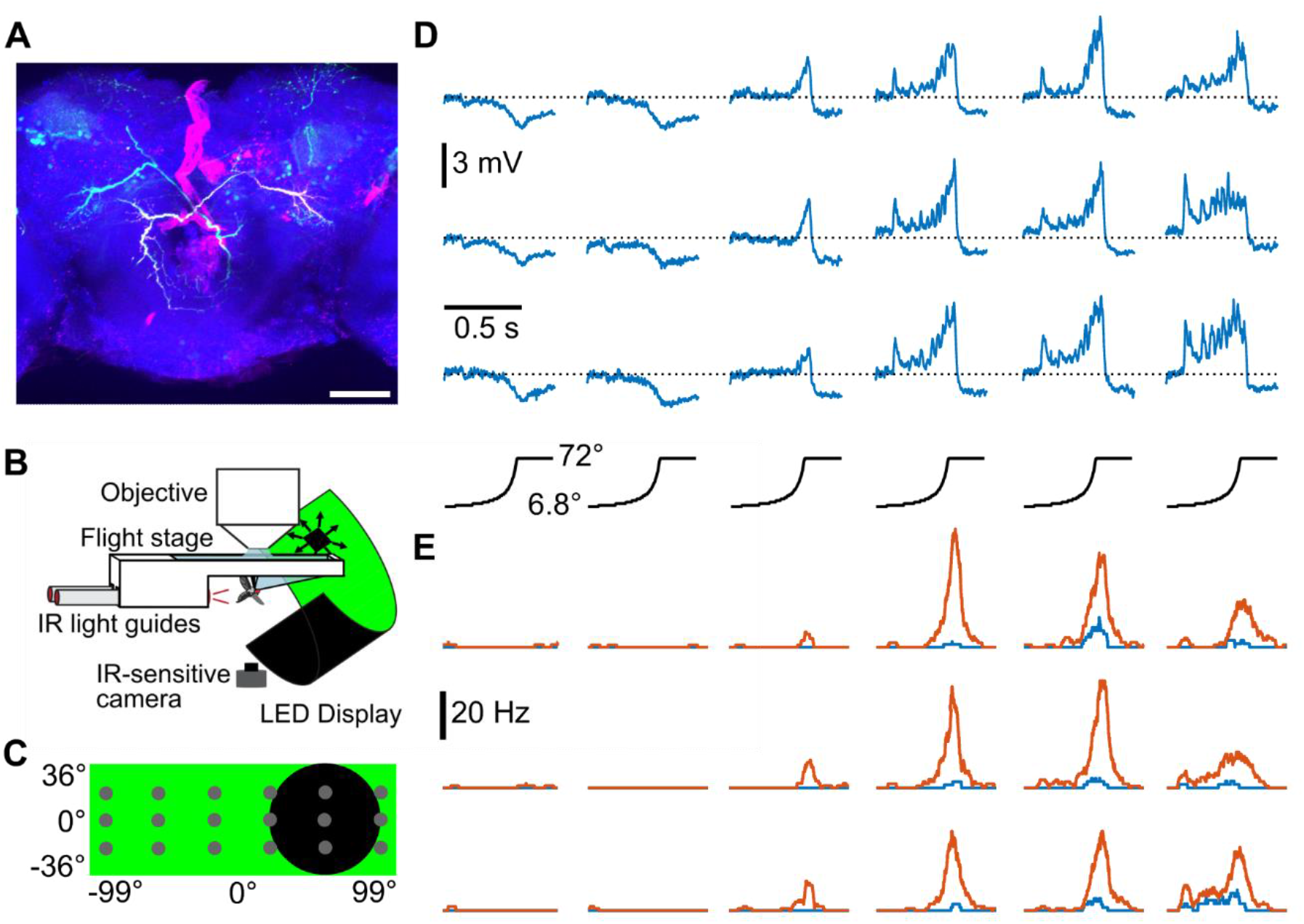
Responses of DNp03 to looming stimuli. **A)** Z-projection of GFP expression in split-Gal4-driver line labelling DNp03 (green: anti-GFP) with filled cell shown in white (streptavidin-Alexa568) and anti-NC82 in blue. Scale bar: 50µm. **B)** Scheme of the setup (adapted from [44]). **C)** Scheme of the LED panels with the positions of the looming stimuli used in D and E indicated by gray dots. One stimulus is shown fully expanded (black). **D)** Mean membrane potential changes of DNp03 during looming stimuli (angular size indicated by black traces) centered at different positions (compare C) during rest (blue). N = 10 flies. Shown are the means across flies. **E)** Same as D, but showing the average spike rate of DNp03 during either rest (blue) or flight (orange). N = 6 flies.

To assess visual responses, we presented looming stimuli on our LED panels. These stimuli were centered at 18 different positions (Fig. 1C) and expanded to a maximal size of approximately 72° in diameter within less than 500 ms. When the flies where resting (meaning not flying), DNp03 responded to looming stimuli on the side ipsilateral to its cell body and dendritic tree with a graded depolarization of its membrane potential irrespective of the exact position (Fig. 1D, note that stimuli at the edges of the panels were cut off to some degree). There was a slight inhibition after the end of stimulus presentation. In contrast, looming stimuli presented on the contralateral side lead to a hyperpolarization of DNp03. When the same stimuli were presented during flight, DNp03 reliably fired action potentials in response to looming on the ipsilateral side, which only rarely happened during resting (Fig. 1E, 2A). Therefore, responses of DNp03 are dependent on the behavioral state of the fly.

**Figure 2.**
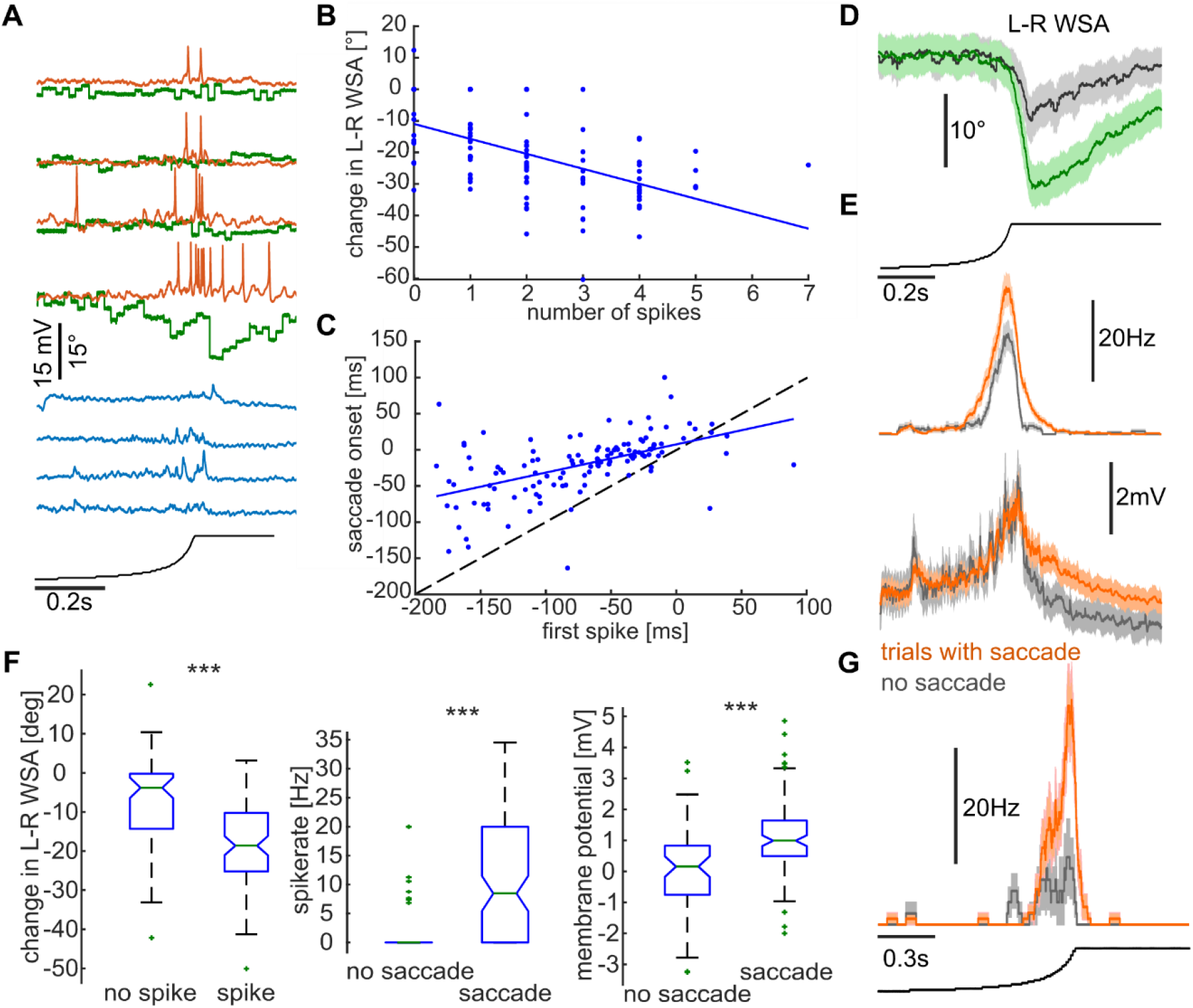
Correlation of DNp03 activity with behavior. **A)** All trials from one recorded neuron to the same ipsilateral looming stimulus during rest (blue) or flight (orange). Corresponding changes in L-R WSA are shown in green. **B)** Number of DNp03 spikes plotted against amplitude of L-R WSA change for one recorded cell (blue dots) for looming stimuli presented on the ipsilateral side (same recordings as shown in Fig. 1). The linear regression is shown as solid blue line. **C)** Saccade onset plotted against time of first spike in DNp03 (blue dots) for looming stimuli presented on the ipsilateral side for all flies (N = 6). The linear regression is shown as blue line. Zero represents the time of collision. The dashed line represents the line of equality. **D)** Mean (+SEM) changes in L-R WSA across all trials, where DNp03 fired either no (dark gray, n = 70 trials) or at least one spike (green, n = 92 trials) after the end of an ipsilateral looming stimulus. N = 6 flies. **E)** Mean (+SEM) spike rate (top) and membrane potential (bottom) across all trials, with (orange, n = 106) or without (light gray, n = 56) a turning response to the ipsilateral looming stimulus. **F)** Statistical comparisons for data shown in E. Box plots represent the baseline-subtracted means in a time interval after stimulus presentation for the different traces (see Methods). p = 6^*^10^-9^ (left plot), p = 8^*^10^-8^ (middle), p = 7^*^10^-9^ (right plot). **G)** Same as E, but for a slower looming stimulus presented at one ipsilateral position. N = 4 flies.

We further characterized DNp03’s response properties with additional visual stimuli. Looming stimuli presented at half the original speed elicited responses that were slightly smaller in amplitude but qualitatively similar to those observed with faster stimuli (Suppl. Fig. 1A). To control for the overall changes in light intensity, we presented a stimulus in which the light intensity was dimmed at the same rate as the looming stimulus, but at random positions within the same area. This control led to a much weaker depolarization in DNp03 (Suppl. Fig. 1B), indicating that the neuron’s response is specific to the expanding nature of the looming stimulus rather than overall luminance changes. Finally, we tested DNp03’s response to a small square moving horizontally at different elevations. This stimulus led to a depolarization only when presented in the most lateral part of our LED panels (Suppl. Fig. 1C), suggesting that DNp03 has distinct receptive fields for looming stimuli and small moving objects. Altogether, these properties - activation by looming stimuli from one side and a stronger response during flight - suggest that DNp03 could mediate a directional turning response away from a potential threat.

### Correlation of DNp03 activity with turning behavior

Generally, we observed variability in behavioral output measured as changes in L-R WSA and the number of spikes DNp03 fired both within the same fly (Fig. 2A) and across flies. Using the data presented above, we therefore tested, whether there was any correlation between activity of DNp03 and the execution and magnitude of a turning response. We call changes in L-R WSA saccades in the following even though flies cannot actually turn in our preparation. As individual flies responded to the looming stimulus either most of the time or almost never, we compared trials across all flies, for which we obtained data during flight. Looming stimuli presented on the ipsilateral side on average led to behavioral responses of similar magnitude across different positions, except for the ones in the corners, which were cut off at two sides (Suppl. Fig. 2A). We therefore pooled trials across all ipsilateral positions except for the two corner positions. We detected saccades manually as a clear deviation of L-R WSA from the noisy baseline (Suppl. Fig 2B), blinded to the neuronal activity during that trial, and determined both the time of onset and peak of the saccade. We first tested, whether there was any correlation between the number of spikes and the amplitude of the change in L-R WSA. A decrease in L-R WSA corresponds to a turn to the left and away from the looming stimulus presented on the right side. For this analysis, we just used data from one fly from which we obtained the most flight trials, because the typical number of spikes fired varied across flies. When plotting the number of spikes against the change in L-R WSA, we found that there was a correlation, but it was generally weak (Fig. 2B, R^2^ = 0.27). Even in some trials, where no action potentials were fired, the fly exhibited a change in L-R WSA, which we interpreted as a saccade.

Using all the trials, in which we detected a saccade from all flies, we looked at the timing between neuronal activity and behavioral onset (see also Suppl.Fig. 2B). Plotting the time of the first spike within a window surrounding the maximal extent of the looming stimulus and saccade onset, which we determined manually, revealed a correlation between the measurements (Fig. 2C, R^2^ = 0.29), with the first spike usually preceding the change in L-R WSA. This does not necessarily imply that the first spike triggers the saccade, as DNp03 usually fired more than one spike in response to the looming stimulus, but only that the earlier the cell was active, the earlier the saccade was usually elicited.

In our preparation, a saccade is usually elicited around the time when the looming stimulus reaches its maximal extent of around 70 deg. While DNp03 is inhibited after this time point during rest, we noticed that in flight trials, DNp03 sometimes continued spiking after the end of stimulus presentation. To explore this further, for each trial we first counted the number of spikes in a time window after the looming stimulus had stopped expanding (see Suppl. Fig. 2B). We then calculated averages of the L-R WSA response for all trials in which there was at least one spike in that time window and for all trials with no spike during that time. In trials with a prolonged spiking response, changes in L-R WSA were significantly larger and lasted longer (Fig. 2D, F). We then did the inverse and grouped all flight trials based on whether a saccade occurred or not and averaged both the spike rate and the raw membrane potential for both groups. While in trials with no saccade, both the membrane potential and the spike rate drop to baseline quickly after the end of the stimulus, in trials with a saccade neuronal activity was sustained (Fig. 2E, F). This was true for both membrane depolarization and spike rate. We observed a similar trend in the data of individual flies and for the slower stimulus that was presented at only one ipsilateral position, although here the spike rate generally reached much higher amplitudes in trials with a saccade even during stimulus presentation (Fig. 2G). These findings suggest that a sustained activity in DNp03 even after the end of the visual stimulus reflects neuronal processes that determine, whether a saccade is initiated or not.

### Activity of DNp03 during spontaneous saccades

We observed that DNp03 exhibited short periods of hyperpolarization or spikes even in the absence of any visual stimulus or during stimuli that did not normally activate DNp03 (Fig. 3A). Often this change in activity coincided with a change in L-R WSA, which we consider a spontaneous saccade. The sign of the change corresponded to what we have observed during looming stimuli, i.e. a hyperpolarization of DNp03 was observed during increases in L-R WSA (rightward turns) and a depolarization or spikes was observed during decreases in L-R WSA (leftward turns). To quantify this effect, we selected times from our data, where no stimulus was shown or where the stimulus remained stationary (in between stimulus presentations). We then detected spikes occurring during those times and averaged the L-R WSA around those time points. If DNp03 fired multiple action potential with short intervals, only the last spike was selected. In this spontaneous spike-triggered average there is a clear decrease in L-R WSA (Fig. 3B) suggesting that spontaneous saccades are also associated with activity changes in DNp03. Together with the fact that the previously described DNaX is also active during both spontaneous and looming-elicited saccades ^20^, this suggest that there is at least cross-talk between the pathways underlying both types of turns.

**Figure 3.**
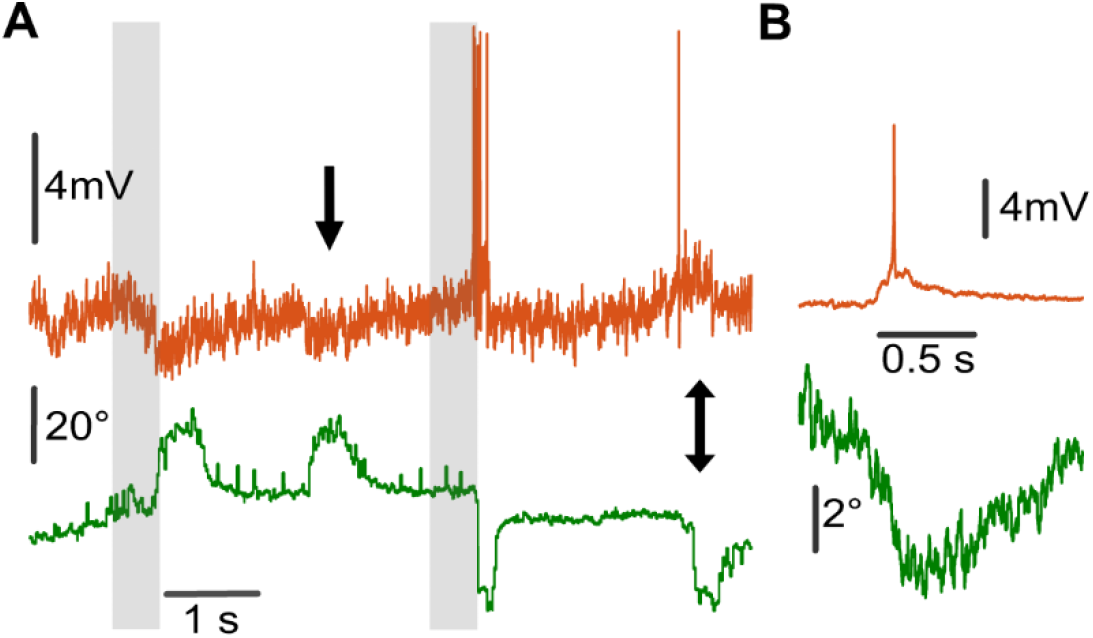
DNp03 activity during spontaneous saccades. **A)** Raw membrane potential (orange) and L-R WSA (green) trace of one DNp03 neuron. Gray shaded areas represent times of looming stimulus presentation from the contra- and ipsilateral side. Arrows indicate spontaneous saccades to the left and right. **B)** Average changes in L-R WSA triggered on spikes detected in between presentation of looming stimuli. N = 6 flies, n = 130 trials.

### Optogenetic activation of descending neurons during free flight

Our previous findings suggest that DNp03 is involved in the initiation and control of flight saccades. However, DNp03 activity alone cannot fully account for the behavioral variability observed in our preparation. This suggests that other descending neurons, such as the previously described DNaX or potentially undiscovered neurons, likely contribute to saccade initiation and control. In addition, in our head-fixed preparation, flies cannot actually perform a saccade. We instead measure just one parameter, L-R WSA, which could be associated with only part of this more complex behavioral maneuver. A free flight saccade typically consists of a banked turn and thus involves rotations around all three body axes ^12^. We therefore wanted to know, whether activity of a single type of descending neuron can influence saccade initiation during free flight. To test this, we built a cylindrical free flight setup allowing us to track the flies’ position in 3D in real-time (Fig. 4A). We used a specific split-Gal4 driver line to express the red-shifted channelrhodopsin csChrimson ^32^ in DNp03 neurons. In our experimental setup, we activated a red columnar light source in the center of the arena for 300 ms whenever a fly was detected in this region. While control flies did not seemingly react to the light stimulus, bilateral optogenetic activation of DNp03 lead to a strong increase in angular velocity (Fig. 4B) associated with a change in flight direction (Fig. 4C). The turning angles showed no directional bias, an expected outcome given our bilateral activation of DNp03. This lack of bias is consistent with the hypothesis that DNp03 neurons on opposite sides of the brain elicit antagonistic turning responses. We assume that inhibitory connections between the two DNp03 neurons or their downstream pathways assure that only one DNp03 neuron is dominant at any giving time. This would assure that during conflicting stimuli, which would activate both neurons to some degree, a directional turning response is nevertheless initiated. Similar results albeit with a slightly different time course of angular velocity changes were obtained when we activated DNaX (Fig. 4B,C). Further experiments will be necessary to show, whether unilateral activation of these DNs is sufficient to elicit turns to only one side and whether there are differences in the type of maneuver elicited by activating DNp03 versus DNaX. Nevertheless, our findings provide evidence that activation of a single type of descending neuron can initiate a complete saccade during free flight.

**Figure 4.**
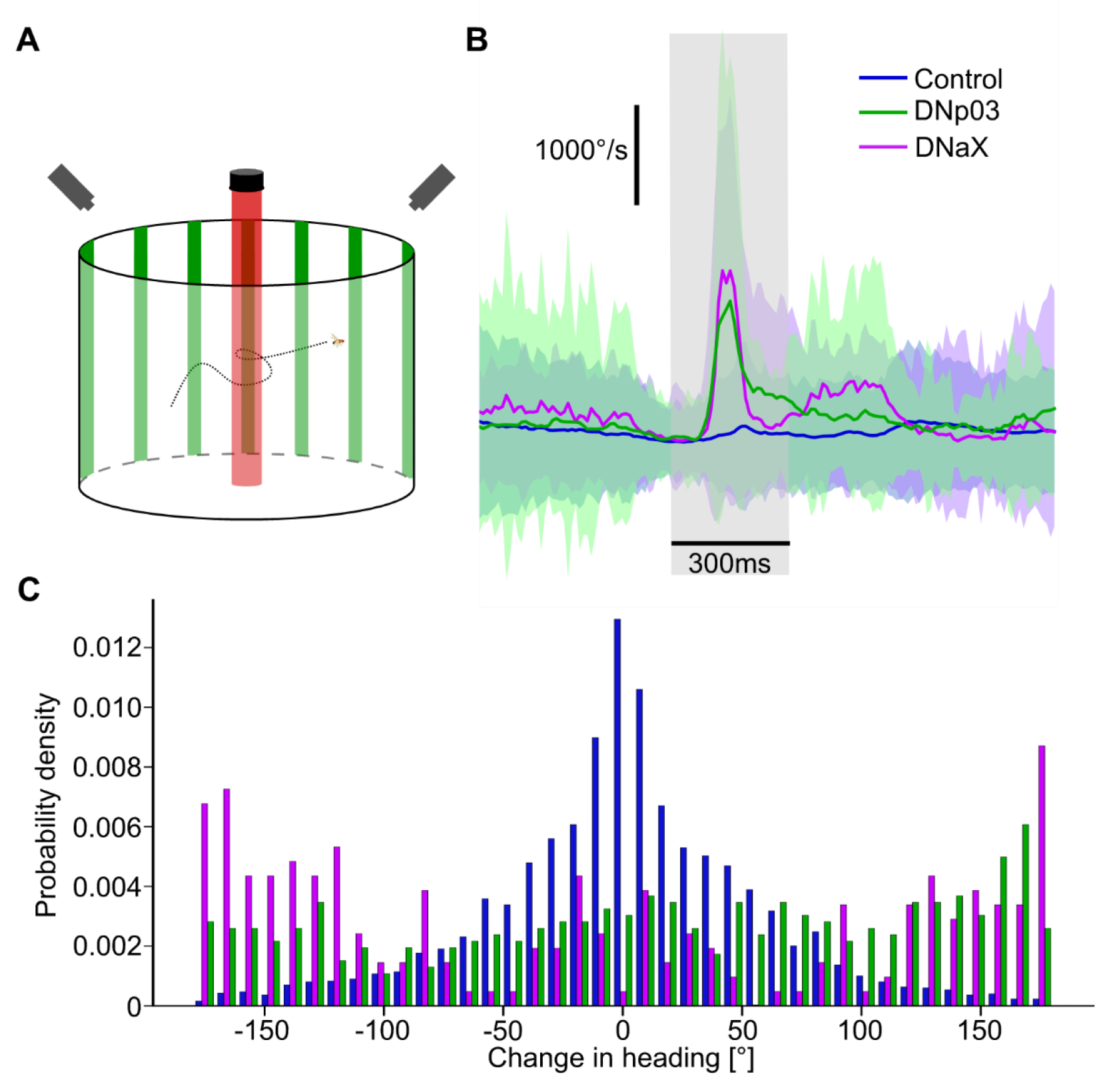
Optogenetic activation of DNp03 and DNaX during free flight. **A)** Scheme of the setup for free flight analysis. Only two of the six cameras are shown for simplicity. **B)** Mean changes (+SD) in angular velocity upon application of the red activation light (shaded gray area) for flies expressing csChrimson bilaterally in DNp03 (green, N = 17 flies, n = 500 traces), DNaX (purple, N = 20 flies, n = 1525 traces) or using empty-split-Gal4 (blue, N = 30 flies, n = 391 traces). **C)** Histogram of the changes in direction upon csChrimson activation for the same flies as in B.

## Discussion

Here we show that the descending neuron DNp03 is responsive to looming stimuli mimicking an approaching object as expected from its connectivity with visual projection neurons. While connectivity with looming-sensitive LPLC1 neurons suggested that DNp03 responses might be biased to stimuli presented ventro-laterally ^26^, we observed relatively uniform responses across stimuli presented on the ipsilateral side. This, however, could be a limitation of the restricted size of our panels, which only covered about 70° in elevation. In addition, several features of DNp03 activity were not predicted by the EM data. First, we showed that contralateral stimuli inhibit DNp03, which is not the case for other descending neurons responsive to looming, such as the giant fiber or DNp06 ^29^. Those were shown to respond to stimuli from both the ipsi- and the contralateral side. Second, its activity is dependent on behavioral state in that it only fires action potential during flight, but not rest, a feature that appears to be common among descending neurons ^25,31,33^. Third, DNp03 also shows changes in activity associated with spontaneous saccades, similar to DNaX. Several DNa neurons are postsynaptic to DNp03 on its contralateral side ^22^. The descending neuron DNaX has been shown previously to be active during both looming-elicited and spontaneous saccades ^20,21^. While the exact cell type in the EM dataset to which DNaX corresponds is still unclear, it seems likely that DNp03 could mediate its activity during looming-elicited saccades.

We furthermore showed that optogenetic activation of DNp03 is sufficient to elicit saccades in free flight, suggesting an important role for this neuron in controlling rapid flight maneuvers. This is supported by the connectivity of DNp03 in the ventral nerve cord, where it makes direct inputs to motor neurons controlling the flight power muscles as well as both inputs directly or indirectly to motor neurons controlling the wing steering muscles, such as i1, b3 and ps1 ^30^. Both i1 and b3 are associated with decreases in wing stroke amplitude ^34–37^. Via interneurons, DNp03 also connects to steering muscles on the opposite side (e.g. b1 and b2) that are associated with increases in wing stroke amplitude and could in this way control an evasive turning response.

Our analysis suggests that there is a small correlation between DNp03 activity and the magnitude of the turning response, measured as change in L-R WSA. A free flight saccade is a complex maneuver, consisting of a banked turn during which flies rotate around all three body axes ^12,38,39^. They first perform a quick roll and counter-roll to change direction and only turn around the yaw (vertical) axis more slowly to realign their body with their new heading. It remains unclear, whether the changes in L-R WSA observed during head-fixed flight, which last longer than a typical free flight saccade, correspond to the roll or the slower yaw maneuver or both. It is therefore possible that DNp03 activity could be more strongly correlated with other aspects of the saccade, which are not captured by our measurement of wing stroke amplitude. We observed occasional changes in L-R WSA in trials, where no spike was detected in DNp03. This could indicate that other descending neurons are involved in initiating the saccade in these cases. Alternatively, subthreshold depolarization of DNp03 could be responsible for eliciting a saccade, as other descending neurons such as DNaX seem to transmit information without action potentials ^20^.

Most strikingly, we observed that in trials, in which the fly performed a saccade, activity in DNp03 – measured as either spike rate or membrane depolarization – was often sustained for up to 100ms even after the end of the looming stimulus and after the time of collision. This sustained activity was unexpected, as DNp03 is typically inhibited immediately after a looming stimulus in resting flies. One possibility of how sustained activity in DNp03 is achieved could involve the suppression of an inhibitory signal that normally terminates the looming response. This suppression, potentially mediated by an unknown pathway, might then trigger the initiation of a saccade. Alternatively, there could be a recurrent excitatory feedback onto DNp03, which leads to the prolonged activity. This explanation also seems plausible, given that recurrent connectivity is very prominent in the *Drosophila* connectome ^40^. The fact that DNp03 activity is sustained even after the presumed time of collision appears surprising. However, it is possible that DN activation is required throughout the duration of the saccade and could mediate e.g. the slower yaw turn. Also, the duration of a saccade is usually longer in tethered compared to free flight, which could be reflected in the longer activation of DNp03.

None of the analyzed parameters of DNp03 activity could explain the full variability of the observed behavior. Therefore, we assume that other descending neurons contribute to the initiation of a saccade. These might me the previously described DNaX, DNb01, DNp06, but likely also other descending neurons responsive to looming that have not yet been described physiologically ^21,27,28^. The direction of escape saccades depends on where the looming stimulus is shown ^12^. Multiple descending neurons might, for example, be tuned to different directions of looming and in concert mediate parameters such as the direction and speed of the saccade. A recent paper showed that descending neurons involved in walking are often strongly interconnected and could thus act as a population even if only one of them is initially activated ^41^. Given that DNp03 similarly makes synapses with other descending neurons, this is likely the case for the control of flight behavior as well. Therefore, even though optogenetic activation of DNp03 elicited saccades in free flight, we cannot conclude that DNp03 necessarily acts as a command-like neuron, as it will activate other descending neurons, which probably act in concert to control the behavior.

While the pathways for controlling spontaneous and escape saccades seem to strongly interact, as both DNaX and DNp03 are active during both types of maneuvers, we still expect them to not be identical. Spontaneous saccades during free flight show important differences from escape saccades in that they are more stereotyped ^38,42^. In contrast, during escape saccades, flies sacrifice flight stability for speed. Therefore, either the pathways controlling these types of manuvers do not completely overlap or there could be differences in the timing of when different pathways controlling different aspects of the saccade, such as the initial roll and the subsequent yaw turn are activated. Spike timing in the giant fiber mediating escape jumps, for example, has been shown to be important for action selection by determining whether a fast, less well-controlled take-off is initiated or a slower more controlled maneuver ^6^.

Further experiments will be necessary to understand how individual descending neurons contribute to finer aspects of the control of flight saccades. Our work provides an entry point into studying the neuronal control of this important behavior and into how behavioral variability is generated by the nervous system, which could be relevant for other systems as well.

## Methods

### Data availability

Data will be made available upon request.

### Flies

Strains of *Drosophila melanogaster* were bred on premixed cornmeal-agar medium (JazzMix *Drosophila* food, Fisherbrand) at 25°C on a 12 h day/12 h night cycle.

For labelling DNp03 for electrophysiology experiments the split-Gal4-line SS01081 (BDSC_75817) ^27^ was crossed to +;+;P{10xUAS-IVS-Syn21-GFP-p10}attP2 (kindly provided by the lab of Prof. Michael Dickinson). For optogenetic activation during free flight w[1118]; P{y[+t7.7] w[+mC]=20xUAS-CsChrimson.mCherry}su(Hw)attP5; + (BDSC_82181) was crossed to the DNp03-line described above, to the split-Gal4-line VT025718.AD; R56G08.DBD ^21^ labelling DNaX or to the empty-split Gal4 w[1118]; P{y[+t7.7] w[+mC]=p65.AD.Uw}attP40; P{y[+t7.7] w[+mC]=GAL4.DBD.Uw}attP2 (BDSC_ 79603) line as a control.

### Electrophysiology and tethered flight behavior

Whole-cell patch-clamp recordings during head-fixed flight were performed as described previously ^43^. Briefly, 2-5 day old female flies were glued to custom-made holders under cold-anaesthesia using UV-cured glue. Legs were removed and a small whole was cut into the cuticle above the DNp03 cell bodies. The neural sheath above the cell bodies was removed using a pipette filled with collagenase and pressure application. Recording electrodes filled with intracellular solution containing Biocytin and Alexa568 had a resistance of 6-9 MOhm. The extracellular solution was oxygenated with Carbogen before the experiment, but no perfusion was done during experiments, which does not seem to affect the quality of recordings. Only cells with a stable membrane potential of less than -35mV were included in the analysis. In total, data from 12 recorded neurons were included in the analysis and for six cells of these we obtained recordings during flight. For five of the cells we obtained data for the slower and the faster looming stimulus. For three of them, we also measured responses to the control stimulus and to the moving square.

L-R WSA data were recorded using Kinefly at a frame rate of 50 Hz ^31,43^. Those data were not filtered, which is why individual traces do not look smooth. No corrections were made for measurement delays, which can be on the order of 30ms ^44^.

### Immunohistochemistry

Antibody staining were performed on whole-mount *Drosophila* brains according to established protocols. The following antibodies were used to visualize cells after intracellular recording: anti-GFP polyclonal (1:1000, rabbit, ThermoFisher Scientific, MA, USA, LOT# 2477546) primaryantibody, the polyclonal secondary antibody anti-rabbit AlexaFluor488 (1:200, goat, ThermoFisher Scientific, LOT# 1966932), Strepavidin-AlexaFluor568 (1:1000, ThermoFisher Scientific) to stain against Biocytin, which was delivered via the patch pipette.

### Visual stimuli

Visual stimuli presented on green LED panels as described previously ^43,45^. The panels covered about 198° along the azimuth and 72° in height with each pixel covering about 2.25 degrees.

The looming stimuli consisted of dark circles on a bright background mimicking an object approaching at constant speed. They expanded from a perceived diameter of about 6.75° to its maximal extent of about 72° and then remained stationary. The half-size to approach ratio l/v was 30ms for the faster looming stimulus and 60ms for the slower stimulus. For the control stimulus, random pixels within the circular area covering the maximal extent of the looming stimulus turned dark such that the light intensity in this area decreased at a similar time course as for the slower looming stimulus. The faster looming stimulus was presented at 18 different positions representing combinations of 6 different positions along the azimuth (±90°, ±54°, ±18°) and 3 different elevations (-18°, 0°, +18°). The slower stimlulus was presented at three different positions along the azimuth (frontal and ±54°).

To test for responses to small moving objects, we presented a 4×4 pixel black square on a bright background moving from left to right and back again at eight different elevations in a randomized order. The speed was 66°/s.

### Electrophysiology data analysis

All data obtained in the tethered preparation were analyzed using Matlab (Mathworks). Analysis scripts are provided upon request. Trials in which the visual stimulus did not follow the proper time course were discarded.

#### Spike detection and calculation of spike rate

The raw membrane potential data were filtered with a 6^th^ order bandpass filter with cut-off frequencies of 5 and 300 Hz. In the filtered signal, we detected positive peaks surpassing a threshold that was empirically determined for each recorded cell. We then selected the time points, at which the unfiltered membrane potential reached a maximum, within a 7ms interval around the detected peaks. If the membrane potential surpassed a second threshold, this point was considered to be the time of a spike. We visually verified proper detection for each recording. We choose the threshold such that smaller spike-like events that also occurred during rest, which were clearly smaller in amplitude and distinct from the spikes we observed during flight, were not detected by our algorithm. To estimate spike rate in each trial, we counted the number of spikes in a sliding window of 50ms length and divided the obtained number by this duration. We chose this time interval, as it resulted in relatively smooth curves while preserving sufficient timing information.

#### Statistics

No statistical tests were used to predetermine sample sizes, which were, however, in the range of other publications in the field. Detailed numbers including ‘N’ of individual animals and ‘n’ of trials are given in every figure legend.

The maximal change in L-R WSA (Fig. 2B) was determined by averaging L-R WSA in a time interval of 50 ms after the manually determined peak of the saccade and the baseline calculated as the average L-R WSA in the first 100 ms of stimulus presentation was subtracted from this value. The number of spikes were counted in a time interval from 0.4 to 0.6 seconds after start of stimulus presentation.

For the analysis shown in Fig. 2C, the first spike was selected in a time interval of 0.3 to 0.8 s after start of stimulus presentation. Trials with no spikes and no saccades were discarded for this analysis.

For the analysis shown in Fig. 2D, the number of spikes were counted in a time interval from 0.5 to 0.6 s after start of stimulus presentation and trials were separated based on whether the spike count was zero or at least one.

For Figs. 2E-G, trials were divided into groups based on the criteria mentioned in the Results. Statistical comparisons between groups were done using a Wilcoxon rank sum test performed on baseline-subtracted averages of the following time intervals: 0.52 to 0.62s for L-R WSA, 0.51 to 0.54s for spike rates and 0.5 to 0.6s for membrane potential changes. Results were robust against the exact duration of these time intervals.

### Optogenetic activation during free flight

#### Flies

The flies for optogenetic stimuli experiments were reared on a standard media until 3-5d post-eclosion. Then, the flies were transferred to ATR-supplemented and light-shielded food vials containing 25ul of 100mM ATR (R2500, Sigma-Aldrich). The flies were maintained on the ATR media for at least 24 hours before experiments. Directly before the experiments, flies were anesthetized inside the vial at 4°C and then directly transferred to the experimental arena. Recordings of the flies lasted until enough traces were collected, up to 24 hours.

#### Behavioral Arena

The free-flight arena consisted of a custom-built cylindrical plexiglass tube (30cm height, 50cm diameter) with a white diffuser bottom (LED Weiss WH14 GT, Plexiglas, Germany) and a removable top, housed in a room maintained at 18°C. While running experiments, a controlled humidifier (ReptiZoo Reptile Humidifier, Repti-Zoo, FL, USA) was connected to the arena to maintain the interior at 60% relative humidity. For back-illumination, a custom-built circular array of 27 850nm LEDs (VSMA1085250X02, Vishay Intertechnology, Inc., PN, USA) was placed 5 cm below the bottom of the arena. The arena was surrounded by ten flexible LED panels (Linsn LED, Shenzhen, China), with a white diffuser paper between the panels and the walls. For our experiments, a static image consisting of a random array of black rectangles on a white background was displayed on the screens surrounding the arena.

#### 3D Tracking

Real-time 3D tracking of flies was performed at 100Hz using an array of 6 Basler Cameras (two Basler ace U acA800-510um and four Basler ace 2 Basler ace 2 R a2A1920-51gmBAS, Basler AG, Ahrensburg, Germany) positioned above the arena (Fig. 4A). Each camera was equipped with an IR-pass filter (HOYA R72 INFRARED filter, Kenko Tokina Co., Tokyo, Japan), so tracking was performed using only the infrared light. The tracking was implemented in the open-source software Braid ^46^. The intrinsic calibration was performed using an 8×6-checkerboard pattern with the ROS camera calibration package (*Stanford Artificial Intelligence Laboratory et al. (2018). Robotic Operating System. Retrieved from https://www.ros.org).*

#### Experimental Paradigm

All experimental paradigms were developed using Python, Arduino, and Rust and are available at https://github.com/mpinb/braid-opto-arena.

Our system received the tracked fly’s real-time position, which activated a trigger function when the fly was within a pre-defined “trigger zone” of 5 cm in diameter and 10 cm in elevation within the center of the arena. The optogenetic stimulus was delivered via a Thorlabs collimated LED (M625L4-C1, Thorlabs, USA) mounted directly above the center of the arena, with an average power density of 200 μW/mm^2^ (PM121D, Thorlabs, USA). Each optogenetic light pulse activated the LED at full power continuously for 300ms. Every experiment included a 10-second interval between consecutive activations, in which flies could not trigger another event.

#### Data analysis

All analyses were done using Python 3.12. Only trajectories with an optogenetic stimulus were selected for analysis.

Angular velocity was calculated through a series of computational steps. Initially, the arctangent (atan2) of the y- and x-velocity components was computed for each trajectory. The resulting angular data was then unwrapped to prevent discontinuities. To obtain the angular velocity, the time derivative of the unwrapped data was calculated by computing the difference between consecutive angles and dividing by the sampling interval of 0.01 seconds. A 150-frame window centered on the stimulus initiation point was extracted for subsequent analysis, comprising 50 frames pre-stimulus and 100 frames post-stimulus.

To quantify heading changes in response to the stimulus, we employed a comparative approach examining pre- and post-stimulus flight directions. The pre-stimulus heading was calculated as the average direction of movement between frames 40 and 50 of the extracted trajectory, corresponding to 10-0 frames before stimulus onset. The post-stimulus heading was computed as the average direction of movement between frames 50 and 80, representing 0-30 frames after stimulus onset. The change in heading was then determined by comparing these pre- and post-stimulus directions, providing a measure of the fly’s response to the presented stimulus.

## Author contributions

Conceptualization: E.B., B.S.; Methodology: E.B., B.S.; Formal Analysis: E.B., B.S.; Investigation: E.B., B.S.; Writing (Original Draft): B.S.; Writing (Review & Editing): E.B:, B.S.; Visualization: E.B. B.S.; Funding Acquisition: B.S.; Supervision: B.S.

## Acknowledgements

We thank Michael H. Dickinson for sharing fly stocks. This work was supported by an Emmy-Noether grant from the German Research Foundation (to B.S.), an ERC Starting Grant (MOBY-FLY, to B.S.), and the Ministry of Culture and Science of the State of North Rhine-Westphalia (iBehave).

