## Supplementary Figures for "Sustained activity in a descending neuron is associated with flight saccades in *Drosophila*"

**Supplementary Material**

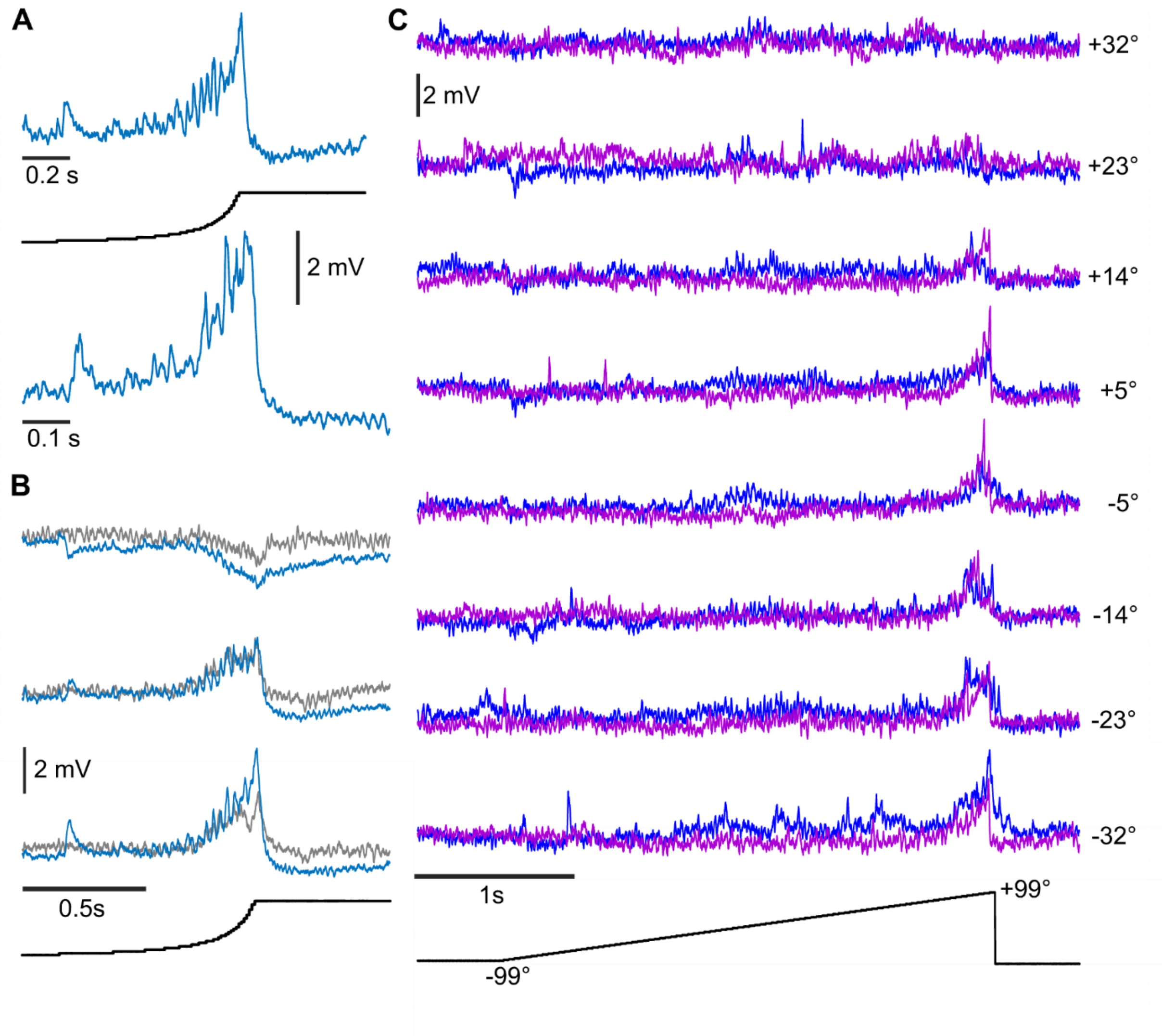


**Supplementary Figure 1. A)** Mean changes in membrane potential for two different velocities of the looming stimulus presented at a single position on the ipsilateral side. The lower plot represents data from Fig. 1. N = 5 flies. The time scales were adjusted such that the change in angular size of the stimulus (black curve) is similar for both traces. **B)** Mean membrane potential changes in response to the slower looming stimulus (blue) and a control stimulus of decreasing light intensity (light gray) presented at a contralateral (top), frontal (middle) and ipsilateral (bottom, data from A) position during rest for the same flies. N = 3 flies. **C)** Mean membrane potential changes in response to a moving square moving from left to right (blue) and right to left (purple) presented at different elevations indicated on the right during rest. The black line represents the position of the square along the azimuth. Traces for leftward motion were inverted in time. N = 3 flies.

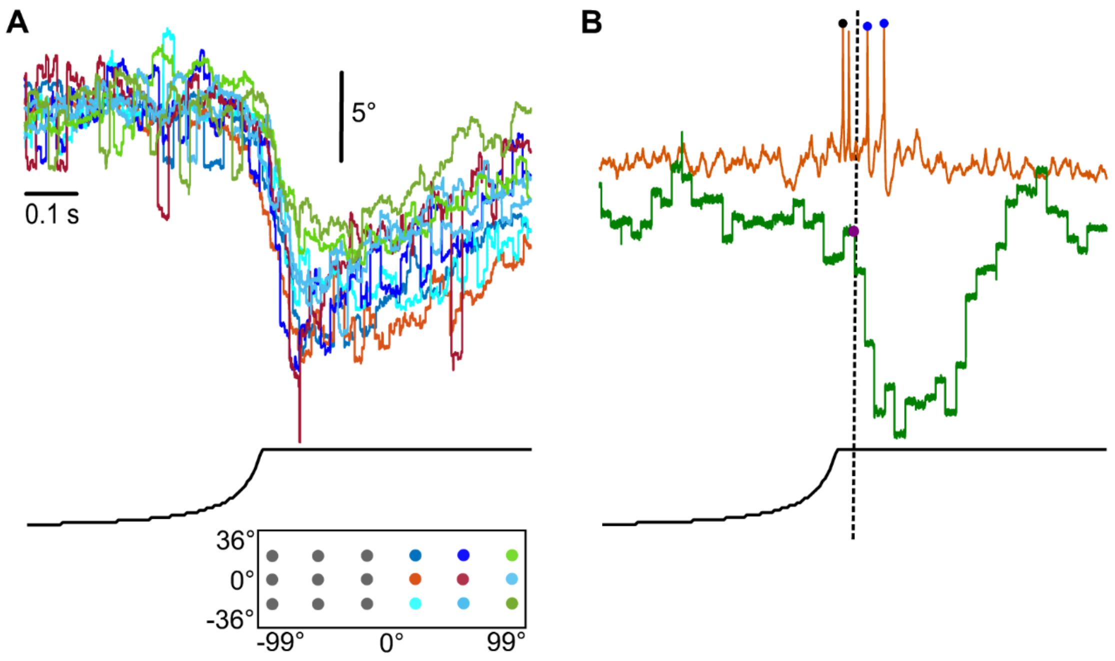


**Supplementary Figure 2. A)** Mean L-R WSA changes during flight obtained during recordings presented in Fig. 1. for all ipsilateral positions of the looming stimulus. Traces are color-coded based on the position of the stimulus as indicated in the inset. **B)** Example trace of DNp03 membrane potential and change in L-R WSA in response to a looming stimulus indicating the first spike (black dot), saccade onset (purple dot) and spikes fired after the time of collision (dotted line, blue dots).
